# Adaptations of an RNA virus to increasing thermal stress

**DOI:** 10.1101/160549

**Authors:** Sonia Singhal, Cierra M. Leon Guerrero, Stella G. Whang, Erin M. McClure, Hannah G. Busch, Benjamin Kerr

## Abstract

Environments can change in incremental fashions, where a shift from one state to another occurs over multiple organismal generations. The *rate* at which the environment changes is expected to influence how and how well populations adapt to the ultimate environment. We used a model system, the lytic RNA bacteriophage Φ6, to investigate this question empirically. We evolved viruses for thermostability by exposing them to heat shocks that increased to a maximum temperature at different rates. We observed increases in the ability of many heat-shocked populations to survive high temperature heat shocks, and on their first exposure to the highest temperature, populations that experienced a gradual increase in temperature had higher average survival than populations that experienced a rapid temperature increase. However, at the end of the experiment, neither the survival of populations at the highest temperature nor the number of mutations per population varied significantly according to the rate of thermal change. We also evaluated mutations from the endpoint populations for their effects on viral thermostability and growth. As expected, some mutations did increase viral thermostability. However, other mutations *decreased* thermostability but increased growth rate, suggesting that benefits of an increased replication rate may have sometimes outweighed the benefits of enhanced thermostability. Our study highlights the importance of considering the effects of multiple selective pressures, even in environments where a single factor is changing.

## Introduction

The process of *de novo* adaptation is typically studied in the context of the simplest form of environmental change, an abrupt shift from an old environmental state to a new one (e.g., [1]). Immediately following the environmental change, the mean fitness of the population shifts to a relatively low value (that is, it is poorly adapted to that environment). The population’s mean fitness then increases as the population gains and fixes beneficial mutations. For instance, under Fisher’s geometric model [2], the population is expected first to fix mutations that confer large gains in fitness, followed by mutations of increasingly smaller beneficial effect as the population approaches the optimal phenotype in the new environment [1].

However, natural environments rarely change in the simple, abrupt fashion assumed by such models. Rather, environmental changes can occur more gradually, on scales that encompass multiple organismal generations. For example, shifts between glacial and interglacial periods occurred over thousands of years (e.g., [3]). Even changes that are rapid on geological scales, such as anthropogenic climate change (e.g., [4]) or changes in pollution levels (e.g., [5]), occur over multiple decades. Adaptation may proceed very differently in such cases of incremental environmental change [6-8].

Evolution in an incrementally changing environment is often modeled as a single quantitative trait evolving under Gaussian stabilizing selection in conditions where the optimal phenotype is constantly shifting [6, 8-13]. In contrast to adaptation under rapid environmental change, adaptation under gradual change is more likely to proceed via fixation of mutations that provide small shifts in phenotype and thus small increases in fitness [6-8]. These shifts allow an evolving population to track the optimal phenotype, but with a lag. The larger the phenotypic lag, the lower the mean fitness of the evolving population. The rate of environmental change can influence the adaptive process by setting the rate at which the population must track changes in the optimal phenotype. More rapid changes in the optimal phenotype typically result in a larger lag of the quantitative trait [6, 8, 11, 12, 14-16]. If a larger distance between the population’s mean phenotype and the optimal phenotype also results in the death of a larger number of individuals, then sufficiently rapid environmental change can lead to population collapse, as small populations lose genetic variation necessary for adaptation [11-13, 17, 18].

The assumptions made by theoretical models may not always be met in biological systems. For this reason, empirical studies using microorganisms have been important in refining our understanding of evolution in incrementally changing environments. In some studies, and in line with model predictions, the rate of population extinction is lower under more gradual environmental change [19-21]. While models predict a higher mean fitness under gradual than rapid environmental change (due to a smaller lag between the population’s mean phenotype and the optimal phenotype), these models tend to consider unlimited change. In contrast, many experiments set limits on the maximum amount of change in the environment. In this framework, the level of environmental stress increases at different rates up to a maximal level, such that treatments involving more rapid change reach the maximum sooner and remain at the maximum longer. Such studies reveal heterogeneity in results of adaptation at the environmental limit. Exposure to low levels of stress can sometimes increase the probability that a population will survive at the environmental limit [20-23]. In some studies, adaptive phenotypes obtained under gradual environmental change have higher fitness in the most stressful environment than phenotypes obtained under rapid environmental change [7, 24, 25]. In other studies, adaptive phenotypes obtained under rapid environmental change are fitter [19, 20, 23]. One study found that the rate of environmental change did not affect fitness in the ultimate environment [26].

Empirical studies also reveal complexities in how the rate of environmental change affects the amount of genetic variation present during the adaptive process. Higher population sizes and less extreme selection coefficients under more gradual environmental change may permit greater genetic diversity [21, 24]. In asexual microbial populations, clonal interference, where distinct beneficial mutations arise in different genetic backgrounds and cannot recombine [27], may also be more prominent under gradual environmental change if multiple mutations of small effect are available simultaneously [24, 25]. On the other hand, when environmental change cannot exceed a maximal value, populations under gradual change must survive a greater range of environments, while populations experiencing the most rapid change must only survive the most extreme environment. If the exposure to a greater diversity of selective environments constrains mutations beneficial in all environments, then a greater diversity of mutations would be predicted under rapid change [7]. Consistent with this hypothesis, some studies find greater variability in phenotypes [7, 19] or fixed mutations [25] under rapid than attenuated environmental change.

Given the heterogeneous results from prior experiments, further experimental studies with different organisms and environmental factors are warranted. In this study, we exposed populations of the lytic RNA bacteriophage Φ6 to heat shocks that increased to a high temperature maximum at varying rates (suddenly, moderately, or gradually; Figs 1 and 2). Subjecting the viruses (but not the host) to heat stress promotes the evolution of thermotolerance via greater stability of viral proteins: Only viruses that survive heat stress with little enough damage that they can subsequently infect a host cell are able to replicate. To track adaptation over time, we measured the percent of the viral population that was able to survive heat shock at each transfer.

**Fig 1:**
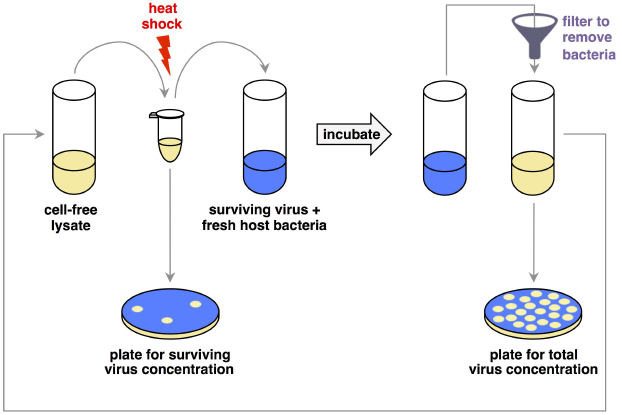
Schematic of the evolution experiment. Bacteria-free lysates of virus were heat shocked for 5 minutes at a pre-specified temperature (Fig 2), then added to culture with naïve host bacteria and grown overnight at 25°C. After the growth period, the viruses were separated from the bacteria by filtration, and the new cell-free lysate was again heat shocked. In order to track changes in survival to thermal stress, viral lysates were plated for concentration before and after heat shock by mixing a dilution of the lysate with abundant host bacteria in soft agar and spreading it onto a Petri dish. After overnight incubation, we counted plaques in the bacterial lawn, each of which originated from a single viral particle. The experiment ran for 32 transfers (approximately 100 viral generations).

**Fig 2:**
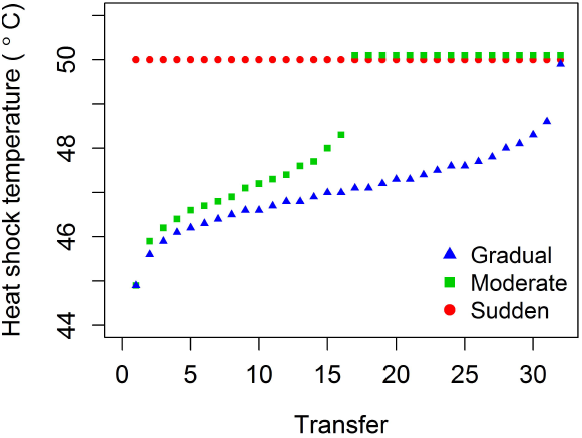
Temperature regimes for experimental evolution with varying rates of thermal change. Points are offset vertically at 50°C for purposes of visualization. Temperature increments were chosen such that the ancestral virus would experience constant (i.e., linear) decreases in its probability of survival. A control regime of heat shocks at a constant 25°C (not shown) accounted for evolutionary change under transfer conditions. See also S2 Table.

We aimed to address how varying rates of thermal change would affect viral evolution in:

*1) Survival on the first exposure to the most extreme environment.* Assuming that mutations that enable survival at intermediate temperatures also contribute to survival at the highest temperature, we predicted that populations under gradual thermal change would have a greater survival when they first reached the highest temperature because they had more time to gain and fix thermostabilizing mutations.

*2) Survival in the most extreme environment at the end of the experiment.* On the one hand, populations that experienced the most rapid change in temperature would also have more time to adapt under the highest temperature, favoring higher final fitness in this treatment. On the other hand, if mutations selected under intermediate temperatures serve as genetic backgrounds for additional mutations that confer high fitness under high temperatures, then populations that experienced a more gradual change in temperature might have higher survival in the ultimate environment (see [21] for a discussion of such epistatic effects in the context of adaptation in changing environments).

*3) The quantity and effect sizes of mutations that permit survival at high temperatures.* We predicted that, in environments that changed rapidly, thermostability would increase more often through single mutations of large effect, while in environments that changed more gradually, thermostability would increase more often through multiple mutations of small effect.

While we did find that evolution at intermediate temperatures enhanced the ability of populations to survive their first exposure to the highest temperature, similar adaptive endpoints were accessible under all rates of environmental change, and the ultimate survival at the highest temperature did not differ across treatments. We also found no strong relationship between the rate of environmental change and the number or effect size of mutations. In addition, we found that selective pressures orthogonal to those of the changing environment can still play a major role in shaping adaptive solutions in stressful environments.

## Methods

### Strains and culture conditions

A list of all viral and bacterial strains used or engineered in this study appears in S1 Table.

□ 6 Cystovirus has a tripartite genome made of double-stranded RNA. The particular strain used in this study originated from three plasmids containing cDNAs of each of the wild type Φ6 segments [29, 30]. These plasmids were co-transformed into bacterial host cells (*Pseudomonas syringae* pathovar *phaseolicola*) to make phage particles [29, 30] (see Reverse engineering, below, for details). L. Mindich (Rutgers University, Newark, New Jersey) kindly provided the following strains: LM4286 (contains pLM687 with the □6 L segment [31]), LM4284 (contains pLM656 with the □6 M segment [32]), LM4285 (contains pLM659 with the □6 S segment [33, 34]), and LM987 (contains pLM857 with the □6 M segment and a lacH marker, which creates phages that make blue plaques [35]).

The laboratory bacterial host for Φ6 growth, *P. phaseolicola* HB10Y, derives from ATCC #21781. Transformation of the phage plasmids was performed into LM2691, a variant of *P. phaseolicola* HB10Y containing a plasmid with a T7 reverse transcriptase [36, 37] (see also Reverse engineering). Both of these hosts were kindly supplied to our laboratory by C. Burch (University of North Carolina, Chapel Hill). During competition assays, counts of clear and blue Φ6 plaques (made from plasmid pLM857) were distinguished on bacterial lawns of a second HB10Y variant, LM1034 (kindly provided by L. Mindich), which contained a plasmid with a lac omega gene [35].

Host cultures were initialized from individual colonies and grown overnight at 25°C in LC medium (Luria-Bertani broth at pH 7.5). Antibiotics (15 μg/mL tetracycline or 200 μg/mL ampicillin) were added to cultures of LM2961 or LM1034, respectively, to maintain their plasmids.

Each viral lysate was prepared from a plaque that had been isolated and stocked in 500 μL of 4:6 (v/v) glycerol:LC. A diluted sample of the virus stock was mixed with 200 μL of stationary-phase of *P. phaseolicola* in LC 0.7% top agar. The mixture was overlaid on an LC 1.5% agar base, and the agar plate was incubated overnight at 25°C. Plaques were collected and filtered in 3 mL of LC medium through cellulose acetate filters (0.2 μm pore, Thermo Scientific) to remove bacterial cells.

### Evolution experiment

The evolution experiment was initialized with a lysate made from a single plaque that had resulted from the transformation from plasmids pLM687, pLM656, and pLM659 (see also Reverse engineering), prepared as described under Strains and culture conditions. This lysate was divided among 20 populations across four treatments with five replicates each (5 Gradual populations, 5 Moderate populations, 5 Sudden populations, 5 Control populations). Cell-free lysates of each population were heat-shocked at a pre-determined temperature (Fig 2, S2 Table), then added to culture with naïve *P. phaseolicola* for overnight growth at 25°C. We performed heat shocks on lysates (i.e., without the bacterial host) so that viral evolution was not affected by host heat shock responses. (We had additionally determined that the bacterial host does not survive temperatures above 45°C.)

Our thermal regimes paralleled the design used in other studies [7, 21, 25]:

1) *Sudden:* First and all subsequent heat shocks were performed at 50°C.
2) *Moderate:* Heat shock temperatures increased from 45°C over the course of evolution, reached 50°C halfway through the experiment, and remained at that temperature thereafter.
3) *Gradual:* Heat shock temperatures increased from 45°C and only reached 50°C on the final transfer.
4) *Control:* Viruses only received a mock “heat shock” at their normal growth temperature (25°C).

The exact rates of increase were determined empirically, with reference to survival of the ancestral genotype across a range of 45-50°C. Each increase in temperature in the Gradual and Moderate lineages represented equally-spaced drops in percent survival for the ancestor. (Further details on calculation of treatment heat shock temperatures are available in the Data Repository.)

#### Preparation for heat shock

Lysates were created from overnight liquid cultures by centrifuging 800 μL of culture at 10,000 rcf through a cellulose acetate spin filter with a 0.2 μm pore (Costar). The lysate’s titer was taken as the mean of duplicate titers on *P. phaseolicola* in agar plates. To control for any density-dependent effects of heat shock on viral survival, we adjusted all lysates by dilution to match the lysate with the lowest titer for that transfer (between 8 × 10^9^ and 2 × 10^10^ plaque-forming units [pfu] /mL). We note that, across treatments, lysate titers fell within less than an order of magnitude of each other, and the treatment to which lowest-titer lysate belonged varied across transfers.

The titer-adjusted lysates were diluted and plated on *P. phaseolicola* for their pre-heat shock concentrations.

#### Heat shock

Heat shocks were then performed on the lysates that had been diluted to the same titer. 50 μL of lysate were aliquoted into PCR strip tubes (one tube per replicate population), placed for 5 minutes on a thermocycler (BioRad, C1000 Thermal Cycler) pre-heated to the appropriate temperature (S2 Table), and then chilled on ice. The heat-shocked lysates were diluted and plated for a count of post-heat shock concentrations.

#### Culturing of surviving phages

Viruses that had survived heat shock were then introduced to bacterial host cells in liquid culture for amplification. Cultures used 4 mL of LC broth and were initialized with a 1/100 dilution of naïve, stationary-phase *P. phaseolicola,* and heat-shocked lysate to a final concentration of approximately 2.5 ×10^3^ viral particles/mL. (These concentrations were approximated based on survivals from the previous transfer, because the exact lysate titers were not known until following day.) We used the same initialization concentration of viruses across treatments to ensure equal mutational opportunities. Cultures were incubated for 24 hours at 25°C with orbital shaking to allow the phages to amplify. The cultures were then prepared for the next round of heat shock as described above.

#### Storage

After each transfer, at least 500 μL of each post-amplification population were mixed with glycerol to a final concentration of 40% and stored long-term at -20°C.

### Reverse engineering

All mutations discussed in this paper were constructed on pLM659, the plasmid containing a cDNA copy of the S segment of □6. Mutations were engineered into this plasmid using the QuikChange II Mutagenesis kit (Agilent) following the manufacturers’ instructions. Primers for each mutation are included in S4 Table. Mutagenized plasmids were stored in *Escherichia coli* XL1-Blue bacteria (included in the QuikChange II Mutagenesis kit), and the mutations of interest were confirmed by Sanger sequencing.

Mutation V109I was engineered using mutagenic PCR and T4 ligation. The plasmid pLM659 was PCR amplified from adjacent, overlapping primers with 5’ phosphorylated ends, one of which contained the mutation of interest, using Phusion polymerase (Thermo Scientific) according to the manufacturer’s instructions. To remove (unmutagenized) template plasmid, the PCR product was digested with DpnI (New England Biolabs) according to the manufacturer’s instructions. Approximately 14 ng of DpnI-digested PCR product were used in a T4 ligation (New England Biolabs) according to the manufacturer’s instructions, and the ligation product was transformed into electrocompetent *E. coli* DH5α, prepared as described for LM2691 below, for storage.

Creating phage particles involved transforming plasmids with each of the Φ6 genomic segments into the bacterium LM2691. We made this strain electrocompetent with the following protocol: A culture of LM2691 was grown to stationary phase, then diluted 1/10 into 50 mL of fresh media and grown to exponential phase. The cells were chilled on ice, then pelleted by centrifugation (6 minutes at 2850 rcf) and washed multiple times with the following resuspensions:

1. 50 mL of ice-cold, sterile water.
2. 15 mL of ice-cold, sterile water.
3. 2 mL of ice-cold 10% glycerol.
4. < 1 mL of ice-cold 20% glycerol (exact volume depended on the number of transformations being performed at the time).

The final suspension of cells was aliquoted into 40-μL volumes for working use.

At least 5 ng of a plasmid containing each Φ6 segment (S, M, and L) were combined with the competent cells (in some cases as much as 100 ng of each plasmid were necessary), incubated on ice for 1 minute, and electroporated on an Eppendorf Eporator in an ice-cold cuvette with a 1-mm gap. The cells were resuspended in 700 μL of SOC medium [38], added to 3 mL of LC 0.7% top agar, plated on LC 1.5% agar plates, and incubated overnight at 25°C. Successful transformations were indicated by viral plaques in the bacterial lawn. At least 6 plaques per genotype were stored for sequence confirmation (see Sequencing viral genotypes).

The ancestral genotype for the evolution experiment resulted from transformation of the original plasmids, pLM687, pLM656, and pLM659. For engineered phage, an engineered version of pLM659 was combined with the original versions of pLM687 and pLM656. A version of Φ6 marked with LacH (used for assaying viral fitness) resulted from transformation of plasmids pLM687, pLM659, and pLM857.

### Sequencing viral genotypes

Sequencing was performed from viral lysates originating from either the stored populations from the evolution experiment or the stored reverse engineered plaques, and made as described in Strains and culture conditions. RNA was extracted from the lysates using the QIAamp Viral RNA Mini Kit (Qiagen) and reverse transcribed into cDNA using SuperScript II reverse transcriptase (Invitrogen), following the manufacturers’ protocols. cDNA samples were PCR amplified with phage-specific primers using touchdown cycling (annealing temperature 65-55°C for 10 cycles, reducing the temperature by 1°C each cycle, followed by 25 cycles with a 55°C annealing temperature). The resulting products were given an ExoSap-IT cleanup (Affymetrix) and Sanger sequenced through GeneWiz. Primers were designed to permit 2× coverage of at least 90% of the genome (excluded sections at the segment ends).

Sequence alignments were performed in Geneious v. 10.0.6 and inspected by eye. The ancestral sequence (GenBank Accession # MF352213-MF352215) was built through alignment against sequence files of the plasmids containing the wild type Φ6 segments (provided by L. Mindich). All other sequences were aligned against the ancestral sequence.

### Assaying viral thermostability

We assessed viral thermostability by exposing lysates to heat shocks across a range of temperatures and measuring the change in viral density under each temperature.

Cell-free lysates of each evaluated genotype were prepared as described under Strains and culture conditions and titered on *P. phaseolicola.* Each block of heat shocks included 3-5 unique genotypes (plaques) and the ancestral genotype. To control for any density-dependent effects of heat shock on viral survival, all lysates were diluted to a titer of 2.17 × 10^8^ viral particles/mL (concentration of the lowest-titer lysate across assay blocks). The diluted lysates were plated on *P. phaseolicola* for pre-heat shock titers and were used for heat shocks.

Lysates were heat shocked over a range of temperatures from 25-54°C. For each temperature tested, three replicate samples containing 50 μL of lysate were heat-shocked for 5 minutes in a pre-heated thermocycler (BioRad, C1000 Thermal Cycler), then chilled on ice.

After heat shock, the lysates were diluted and plated for their survival. Survival of the lysates was calculated as the ratio of post- to pre-heat shock titer multiplied by 100. We estimated viral thermostability across the temperature range using an inverse Hill equation:

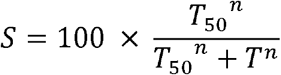

(Equation 1)
where *S* is the percent survival, *T* is the heat shock temperature, *T*_50_ is the temperature with 50% viability, and *n* is the Hill coefficient. A program written in R (version 3.1.2; code available in the Data Repository) estimated parameters *T*_50_ and *n* using maximum log likelihood. Genotypes with a greater *T*_50_ were considered to be more thermostable.

### Assaying viral competitive fitness

Viral growth rates were evaluated through growth competitions against a marked common competitor, the lacH-marked Φ6, under conditions that replicated those of growth during the evolution experiment. Plaques formed by the lacH-marked Φ6 turn blue when plated with X-Gal on LM1034 (a bacterial host containing a plasmid with the complementary lac omega gene), allowing us to distinguish the common competitor from the genotypes engineered for this study.

The common competitor was transformed from plasmids (see Reverse engineering). To pre-adapt this strain to the competition conditions, the plaque isolated from the transformation was passaged for five days in liquid LC medium. However, its growth rate remained low compared to the ancestor of the evolution experiment, so competitions were initialized at a 1:10 ratio of focal strain: common competitor. We confirmed that, for the ancestor, changing its initial ratio in the competition did not affect the measured competitive fitness (S1 Fig).

Lysates of the ancestral virus, each mutant virus, and the lacH-marked common competitor were made up from frozen stocks containing plaques, as described in Strains and culture conditions. Competition mixtures were created by combining the lysate of the focal strain, diluted to 2.89 × 10^8^ pfu/mL, with the lysate of the common competitor, diluted to 2.89 × 10^9^ pfu/mL, in a 50:50 ratio. To obtain initial concentrations of each strain in the competition, the competition mixtures were plated on LM1034 with 100 μL of 40 mg/mL X-Gal (dissolved in DMSO).

Cultures were then initialized from the competition mixtures on the normal *P. phaseolicola* host. Competitions occurred in 4 mL of LC broth with a 1/100 dilution of naïve, stationary-phase *P. phaseolicola.* The competition mixture was added into this culture to a final concentration of approximately 2.5 × 10^3^ viral particles/mL (the initializing concentration used in the evolution experiment; see Culturing of surviving phages, under Experimental evolution). Cultures were incubated for 24 hours at 25°C with orbital shaking.

To obtain final concentrations of each strain, aliquots of the cultures were diluted and plated on LM1034 with 100 μL of 40 mg/mL X-Gal. The competitive fitness of each focal strain was calculated as its change in relative density in the competition over time:

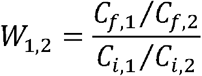

Equation 2)
where *W*_1,2_ denotes the calculated competitive fitness of the focal strain, *C*_*i*_ is the initial concentration, *C*_*f*_ is the final concentration, a subscript 1 denotes the focal strain, and a subscript 2 denotes the common competitor. Relative competitive fitness with respect to the ancestral genotype was then calculated by dividing the competitive fitness of each focal genotype by the mean competitive fitness of the ancestor.

## Results

### Changes in survival to heat shock over time

Viral survival is expected to decrease as the heat shock temperature increases. To account for this effect, we compared the percent survival of populations in the Gradual, Moderate, Sudden, and Control treatments at each transfer to the percent survival of the ancestral genotype at the heat shock temperature experienced in that transfer (Fig 3). If heat-shocked populations did not evolve greater thermostability than the ancestor, then on average there would be no difference between the survival of the population and the ancestor at each temperature (i.e., the point would fall at 0).

**Fig 3.**
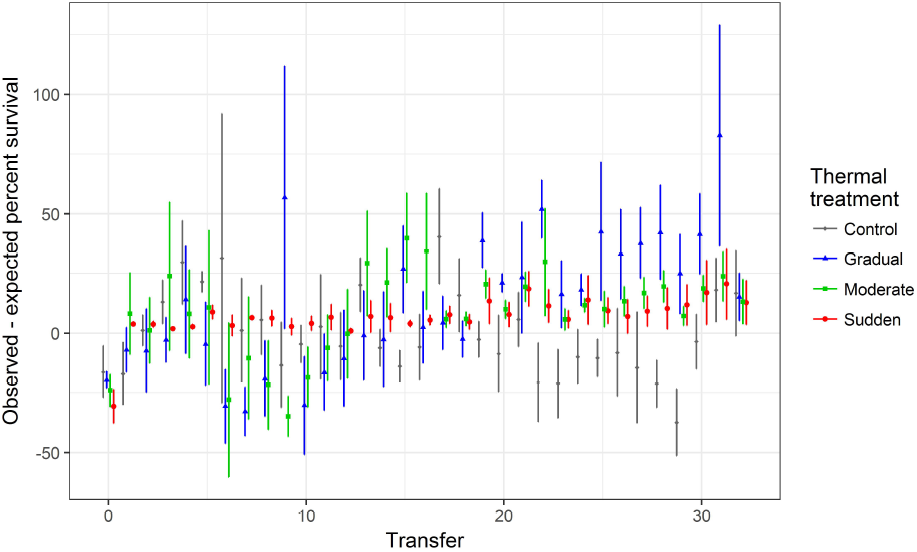
Changes in percent survival of viral lysates to heat shock over time. The survival of each population (Observed percent survival) at each transfer is compared to the percent survival of the ancestor (Expected percent survival) at the temperature used for heat shock (see Fig 2, S2 Table). Points represent the average difference between the population and ancestral survival; error bars represent the standard deviation of this difference. Treatments in which populations evolved better thermostability than the ancestor have a difference greater than 0. Note that, because of stochasticity in determining phage titers, differences occasionally exceed 100% (if more plaques were counted post heat shock than pre heat shock). Points are jittered horizontally for better visualization.

We did not find an improvement over time in the survival of the Control population in response to mock heat shocks at 25°C. In contrast, the percent survival of Φ6 from Gradual, Moderate, and Sudden populations was greater than ancestral values for every transfer in the second half of the experiment (Fig 3). Treatments differed in survival to their respective first exposures to 50°C (analysis of variance, F(2,12) = 4.83, p = 0.03; Fig 4); specifically, populations from the Gradual treatment had a higher survival than populations from the Sudden treatment on their first exposure to 50°C (Tukey’s post-hoc test, p = 0.03; other comparisons were not significant). The survival data thus suggest that heat shocked populations evolved greater thermostability, even during exposure to intermediate temperatures. However, at the end of the experiment, the average survival of Gradual and Moderate populations at 50°C did not differ significantly from survival of populations from the Sudden treatment (analysis of variance, F(2, 12) = 0.0872, p = 0.92; Fig 5).

**Fig 4.**
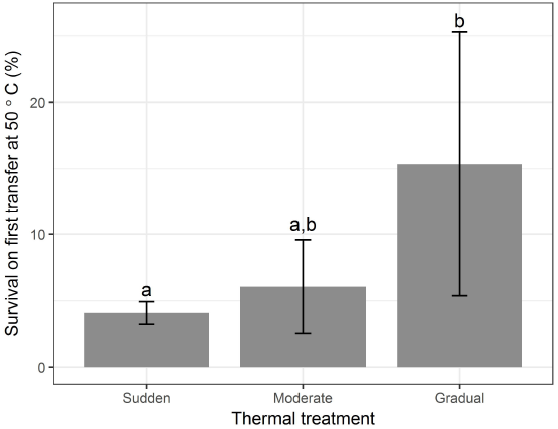
Average percent survival of populations on first transfer at 50°C. Sudden populations first experienced 50°C on Transfer 1, Moderate populations on Transfer 17, and Gradual populations on Transfer 32. Error bars represent the standard deviation of percent survival. Treatments with significantly different percent survivals are denoted with letters.

**Fig 5.**
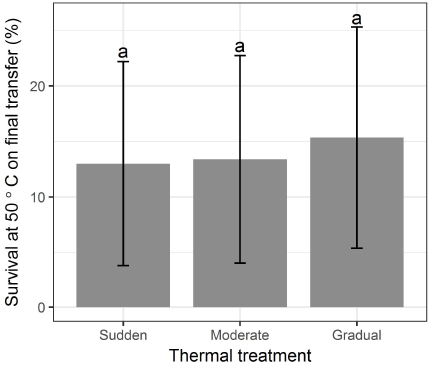
Average percent survival at 50°C on the final transfer (Transfer 32) of the experiment. Error bars represent the standard deviation of percent survival. Percent survival was not significantly different across treatments.

### Genetic basis of thermostability

To identify mutations that may have contributed to increases in thermostability, we sequenced the endpoint populations in gene 5 (encodes the P5 lysis protein) and gene 8 (encodes the P8 outer shell protein). As proteins on the exterior of the virus that are necessary for viral infection [39, 40], both P5 and P8 are expected to experience strong selection for thermostability at high temperatures to maintain their functions.

We found a total of 16 unique mutations across all populations, 11 of which were unique to populations that had experienced high-temperature heat shocks. Although populations from Gradual and Moderate treatments appeared to have accumulated more unique mutations and more mutations per lineage on average than populations from Control and Sudden treatments (Table 1), the number of mutations per population did not vary significantly with treatment (analysis of variance, F(3, 16) = 2.667, p = 0.08).

**Table 1.** Number of mutations in genes encoding for P5 and P8 in each treatment.

| Treatment | Number of different mutations in the treatment | Average number of mutations per population |
| --- | --- | --- |
| Control | 5 | 1.8 |
| Gradual | 7 | 2.4 |
| Moderate | 6 | 2.2 |
| Sudden | 3 | 1.6 |

We reverse engineered 10 of these mutations singly into the ancestral genetic background to evaluate their effect on viral thermostability. Nearly all of the chosen mutations appeared in more than one replicate or had been previously observed in pilot experiments (S3 Table). We evaluated the effects of the single mutations on viral thermostability by exposing bacteria-free lysates of the mutant viruses to heat shocks ranging from 25-55°C, measuring the lysate concentrations before and after heat shock. These data were used to build thermal kill curves, where the percent survivals at each temperature were fit to an inverse Hill equation (Equation 1) using maximum likelihood (Fig 6A).

**Fig 6.**
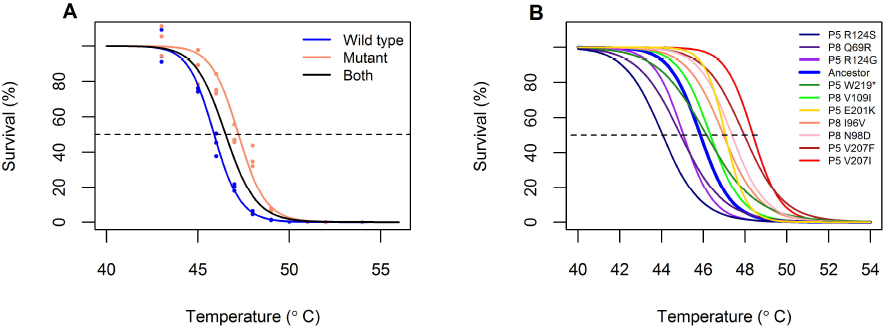
Thermal kill curves of engineered single mutants. A) Calculations of mutant thermostability, using the mutant P8 I96V as an example. Cell-free lysates were exposed to a 5-minute heat shock at each temperature and plated before and afterward to calculate percent survival (circles). (Note that, due to stochasticity in gauging phage titer, phage counts after heat shock can be above phage counts before heat shock, accounting for survivals greater than 100%.) Equation 1 was fitted to the data in R, where the parameters *T*_50_ (intersection of curve with dotted line) and *n* were estimated by maximum likelihood. A first model was fit to the combined data (ancestor + mutant; black curve). A second model then estimated a separate *T*_50_ and *n* for each lysate (blue, ancestor; red, mutant). The latter model was a better fit to the data (log likelihood ratio test, p < 0.0001). B) Empirical thermal kill curves of the ancestor (blue) and ten engineered single mutants, representing the maximum likelihood fit of all measurements taken for each mutant. Data points are omitted for simplicity. In all cases, the model that used a separate *T*_50_ and *n* for the ancestor and the mutants was a better fit to the data (log likelihood ratio test, p < 0.001). Pairwise comparisons with the ancestor, as in part A, can be found in the Data Repository.

The engineered single mutants revealed that different mutations resulted in different gains in thermostability in the ancestral background. As measured by an increase in the *T*_50_ parameter, six of the engineered mutations increased viral thermostability by 0.3-2.1°C while three mutations decreased thermostability by 0.8-1.8°C (Fig 6B). All populations from the evolution experiment for which all mutations were evaluated and that increased in survival had at least one thermostabilizing mutation. (This pattern was most evident in the Sudden treatment; see S2 Fig.)

Four of the six thermostabilizing mutations were conservative mutations for which substitution retained non-polarity of the amino acid, while all mutations that reduced thermostability involved substitutions of polar amino acids to ionically charged amino acids or vice versa. The effect size of the mutations – that is, the amount by which the mutation increased or decreased thermostability with respect to the ancestor – did not differ significantly across heat shock treatments (analysis of variance, F(2, 25) = 0.511, p = 0.61). We note, however, that the number of mutations per population was low and that not all mutations that appeared in each population were evaluated for their effects on thermostability.

#### Growth effects of thermostabilizing mutations

The presence of mutations that *decreased* viral thermostability suggested that these mutations may have fixed because of non-thermal selective pressures. The mutation R124G in P5, for example, appeared in 18 out of 20 different populations, including in the Control treatment. This suggested that the mutation might improve another attribute of fitness, such as viral replication. To test whether destabilizing mutations instead improved replication, we competed all engineered mutants and the ancestor against a common competitor (see Assaying viral competitive fitness in Methods). Many of the mutations appeared to give a competitive growth advantage in comparison to the ancestor, although some mutations decreased viral growth rates (Fig 7). All mutations that reduced thermostability enhanced relative competitive fitness, and many thermostabilizing mutations decreased relative competitive fitness.

**Fig 7.**
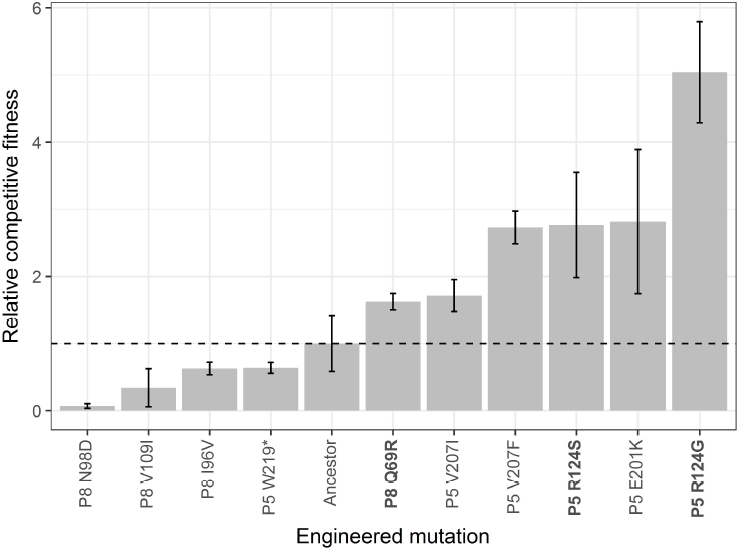
Competitive fitness of engineered single mutants relative to the ancestral genotype. Bar heights indicate the mean of three replicate competitions; error bars denote standard deviation. Mutations in bold font decrease viral thermostability with respect to the ancestor.

A prior study in Φ6 recorded a trade-off between thermostability and growth for one mutation in P5 [41]. In our data set, several individual mutations follow the expected pattern of low *T*_50_ and high growth rate, or high *T*_50_ and low growth rate. To test for a generalized trade-off, we regressed the relative competitive fitness of the mutants against the *T*_50_ values estimated from the thermal kill curves (Fig 8). Although the slope of the regression line was negative, it was not statistically different from a slope of 0 (F-statistic = 0.897, df = 9, p = 0.368).

**Fig 8.**
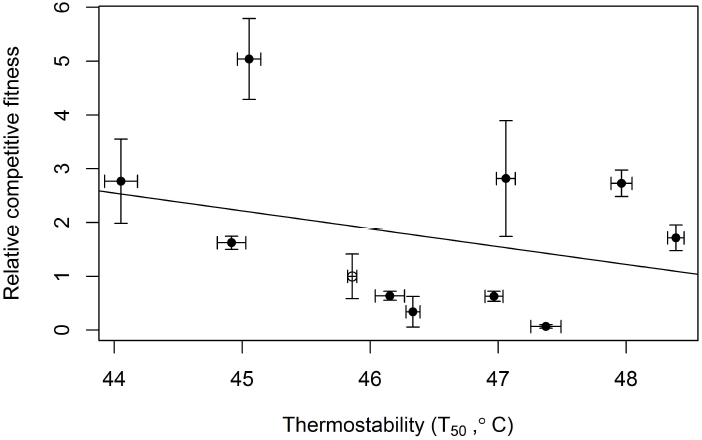
Relationship between relative competitive fitness and *T*_*50*_ for the ancestor and the engineered single mutants. The ancestor is marked with an open circle. X-error bars represent the standard error of the *T*_50_ estimate, while y-error bars represent the standard deviation of three replicate competitions. The line represents the best fit from a linear model.

#### Thermostabilizing and growth effects of combinations of mutations

Several populations from heat-shocked treatments did not appear to have increased their thermostability substantially over time; we expected that mutations from these populations would instead have increased viral growth rates. We confirmed this hypothesis through evaluation of the thermostability and growth rates of genotypes from one of these populations (G1, replicate 1 from the Gradual treatment). Survival of the G1 population had not increased substantially over time; furthermore, it did not carry the common mutation, R124G in P5, that increased viral growth rate and was also found in Control populations (Fig 7, S3 Table). Two genotypes dominated G1 at the end of the experiment. Both genotypes shared the P5 mutation R124S (mutation B in Fig 9), but one genotype also had mutation E201K in P5 (mutation C), while the other had mutation Q69R in P8 (mutation A). As single mutations, all three increased viral growth rates, but only one (E201K) increased thermostability.

**Fig 9.**
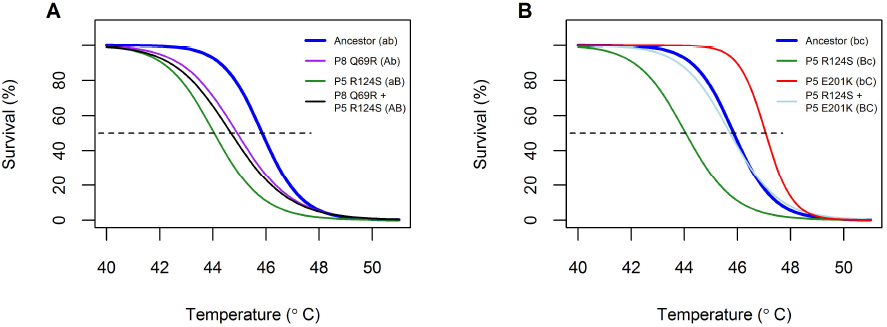
Evaluated thermal kill curves for two combinatorial genotypes found in one Gradual population. Mutations are given secondary labels to denote their allelic state, where a lower case letter (a, b, c) indicates the ancestral residue and an upper case letter (A, B, C) indicates the residue found in the endpoint population. A) Thermal kill curves for double mutant P8 Q69R + P5 R124S and its corresponding single mutants. B) Thermal kill curves for double mutant P5 R124S + P5 E201K and its corresponding single mutants.

We reverse engineered these mutations in their respective double combinations (AB and BC) and evaluated their effects on thermostability and viral growth. Neither double mutant improved thermostability with respect to the ancestor (Fig 9), but both double mutants improved in relative competitive fitness with respect to the ancestor (Fig 10). Interestingly, one of the combinations exhibited sign epistasis for thermostability (mutation P8 Q69R was destabilizing in the ancestral background, but stabilizing in the P5 R124S background).

**Fig 10.**
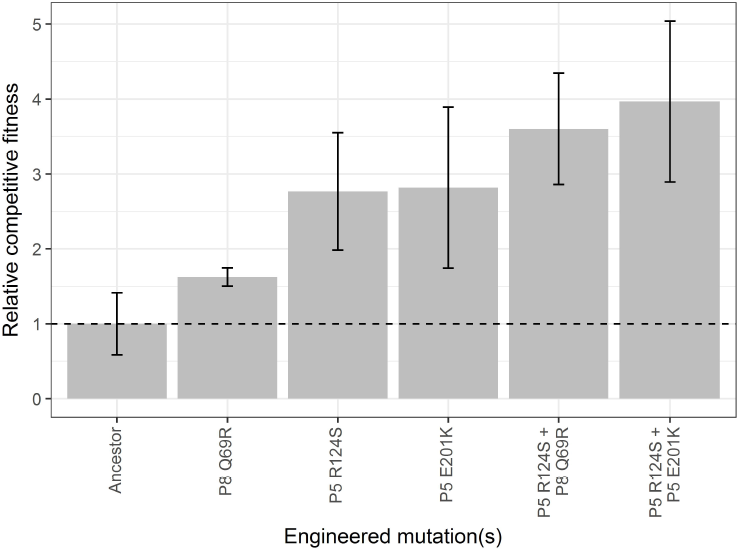
Relative competitive fitness of two combinatorial genotypes found in one Gradual population. Bar heights represent the mean of three replicates; error bars denote standard deviation. The relative competitive fitness of the double mutants is compared to the ancestor and the constituent single mutants. We note that addition of the double mutations did not change the overall relationship between relative competitive fitness and *T*_50_ portrayed in Fig 8.

Based on historical sequencing of the G1 lineage (see S2 Text), the first mutation detected in this population (P5 R124S) decreased thermostability but enhanced viral growth. Subsequent mutations (P8 Q69R and P5 E201K) increased both thermostability and competitive fitness in the presence of P5 R124S. This sequence of mutations is consistent with stronger selection for growth rate early in a gradually changing environment and stronger selection for thermostability later on.

## Discussion

Consistent with prior work in Φ6 [42, 43], we found that virus populations exposed to high-temperature heat shocks evolved greater survival to heat shock, and we identified six causative mutations that increased viral thermostability with respect to the ancestral genotype. We did not find significant differences between Gradual and Sudden treatments in endpoint survival at 5°C, or in the number or effect size of mutations, possibly due to the low number of replicates and mutations in each treatment. Instead, we found that other selective pressures may have been important during the experiment. Specifically, our experimental design permitted two places where selection had a chance to act: on survival, under the high temperature heat shocks; and on replication, when viruses were grown with their bacterial host (Fig 1). Even in heat-shocked populations, we identified mutations that reduced viral thermostability but increased growth rate, suggesting a relatively high selective pressure on viral replication.

Because replication occurred at 25°C, the typical laboratory temperature for Φ6, our experiment is reminiscent of that of Hao *et al.* [44], in which a lytic bacteriophage of *P. fluorescens* was exposed to increasing temperatures punctuated by periods of lower temperature. The authors term the fluctuations to reduced temperatures as periods of “amelioration” because they reduced selective pressures associated with heat stress. If amelioration allowed populations to recover in abundance and *de novo* mutations in the wake of an environmental stress, it could promote adaptation under stressful conditions [22, 45]. On the other hand, because amelioration relaxes the selective pressures present in a stressful environment, it may reduce the likelihood that stress-beneficial mutations will fix [46-49]. Hao *et al.* found that fewer phage populations persisted in treatments that included temperature amelioration than in treatments where the temperature increased monotonically, indicating that periods of amelioration hindered adaptation at high temperatures. Although we did observe increases in thermostability over the course of our evolution experiment, we cannot rule out the possibility that thermostability evolution was hindered due to periods of growth at 25°C.

Amelioration is especially likely to impede adaptation to stressful environments if stress-beneficial mutations impose fitness costs under more benign conditions [44]. A prior study in Φ6 found that a highly thermostabilizing mutation decreased the ability of viruses to replicate at 25°C [41]. Although we did not find support for general trade-off between thermostability and replication in our data (Fig 8), we can identify mutations in P5 and P8 for which we measured high thermostability but low growth rates, or low thermostability but high growth rates. Interestingly, some mutations appeared to increase both thermostability and growth rate. (We note that this last class includes the particular mutation reported in [41], V207F in P5. We speculate that this is because the genetic background of our phage differed from the genotype used in [41].) It is possible that our sample of 11 genotypes is too limited to permit detection of a general trade-off. Alternatively, mutations that contribute to thermostability may not always be constrained by trade-offs. For example, due to their high mutation rates, viruses can find “cost free” adaptations [50-52], which allow them to maintain existing functions while gaining new ones. Such mutations may be particularly important during evolution in changing environments. It is also possible that we did not sample mutations that demonstrate a trade-off, since we examined mutations found at the end of the experiment. Mutations that may have exhibited a trade-off between thermostability and relative fitness and were present at earlier time points might have been outcompeted by mutations that performed well in both dimensions.

That we find growth-enhancing but thermo-destabilizing mutations, however, highlights that organisms experience selective pressures along multiple phenotypic axes. This has potential implications for evolution in environments that change incrementally. For example, a gradually changing thermal environment imposes small differences in selective pressure on the population from generation to generation. In this case, the population may experience stronger relative selective pressure along a non-thermal axis, such as for growth. The population may then not evolve in response to the thermal environment until sufficient thermal change has occurred and relative selective pressures are high enough. For example, Gorter *et al.* [26] report that, in a yeast system, adaptation to general culture conditions preceded adaptation to high metal concentrations under conditions where the metal concentration increased slowly. Similarly, we note that while the first detectable mutation in one Gradual lineage reduced viral thermostability but enhanced growth, both mutations that rose to prominence later in the evolution of this lineage were thermostabilizing in the background of the first mutation (see S2 Text for historical sequencing of this lineage).

In extreme cases, evolution in response to a non-focal selective pressure may impose trade-offs or constraints in the changing stressful environment. Suppose, for example, that evolution for higher growth rates always reduced thermostability. Populations that experienced a gradual increase in temperature may have first fixed growth-enhancing mutations (because of stronger relative selective pressures for growth than for thermostability). However, this would have lowered their thermostability, even as thermal stress became a more prominent selective pressure over time. The population would then be in a *worse* place, in terms of thermostability, than when it started, and mutations of larger thermostabilizing effect would be required to increase its survival at high temperatures. Although we are unaware of any empirical studies that look explicitly at the role of such trade-offs in incrementally changing environments, this conclusion is in the spirit of studies that predict greater phenotypic and genotypic constraint under slow environmental change (e.g., [7, 15, 18, 53]).

Other results from this study are consistent with prior work on adaptation under varying rates of environmental change.

Evidence from prior experiments (e.g., [20-23]) suggests that evolution under mildly stressful environments (such as an intermediate temperature) can enhance a population’s ability to withstand more stressful environments (such as a high temperature). Consistent with this expectation, populations from the Gradual treatment had a higher average survival on their first exposure to 50°C than did populations from the Sudden treatment on their first exposure. In other words, exposure to intermediate temperatures can promote survival of Φ6 at high temperatures.

Several prior experiments find a greater diversity of mutations under rapid than gradual environmental change [7, 19, 25]. In contrast, we find an (insignificant) pattern of more mutations in endpoint Gradual and Moderate populations compared to Sudden populations. This could represent a greater amount of clonal interference in Gradual and Moderate than Sudden lineages (e.g., [24, 28]). (Consider, for example, that lineage G1 had two dominant genotypes at the endpoint of the experiment. Sequencing the lineage at prior time points [see S2 Text] furthermore suggested that both these genotypes were increasing in frequency when the experiment ended.)

Another possibility is that thermostability comprises a set of mutations that vary relatively little regardless of thermal treatment. A study that examined the thermal adaptation of the bacteriophage Qβ found that populations evolved under a constant high temperature did not significantly differ in evolutionary outcomes from populations evolved under fluctuating temperatures [54]. In this study, when we include data from pilot experiments, most of the engineered mutations appeared in populations that had experienced diverse heat shock treatments (S3 Table). Proteins tend to be marginally stable and can be destabilized by a single amino acid substitution [55-57], including by mutations that are adaptive for functions besides stability (e.g., ligand binding [58, 59] or growth [60]). In contrast, computational and empirical data sets suggest that relatively few substitutions will increase a protein’s thermostability [56, 57, 61]. In the case of an enzyme, such as the P5 lysis protein in Φ6, any mutations that increase stability must simultaneously maintain the flexibility or activity necessary for the protein’s function [55]. The number of mutations that increase thermostability may thus be small and/or biochemically constrained for any given protein, resulting in relatively few mutational pathways for improvement.

Overall, our study emphasizes that it is important to take *all* selective pressures into account during an evolution experiment. We found that populations that did not increase in thermostability appeared to have increased instead in replicative ability. We speculate that this may offer an alternative way for populations to persist under heat shock, rather than improving their thermostability: They may be able to make up reductions in population size due to heat shock by increasing their replication rate in its absence. This highlights the conclusion that multiple features of organisms can evolve, even in environments that change in only a single focal factor.

## Acknowledgments

We thank E. Cooper, C. Dunnell, E. Hsieh, and K. van Raay for their assistance in troubleshooting and collecting data for this study; and P.L. Conlin and H. Jordt for comments on the manuscript.

## Supporting information

**S1 Table. Strains used and engineered in this study.** Laboratory collection numbers (BK numbers) are included for request purposes. All other data presented in this study (including this table, other Supporting Information, and the Data Repository) use the project-specific (PRESS) numbers. Mutations are labeled in order ancestral base/amino acid - position - mutated base/amino acid, where the position is measured from the first nucleotide of the NCBI Reference Sequence for the S segment of Φ6 Cystovirus (Accession# NC_003714) or the first amino acid of the protein. We also note the presence of any additional (i.e., non-engineered) mutations present in the mutant viral clones. We account for the effects of these mutations with the “matched” viral clones; see S1 Text.

**S2 Table. Heat shock temperatures used at each transfer in the experimental treatments.**

**S3 Table. Mutations present in genes 5 and 8 at the end of the evolution experiment.** Gene 5 codes for the P5 lysis protein, while gene 8 codes for the P8 outer shell protein. All Gradual (G), Moderate (M), Sudden (S), and Control (C) populations were evaluated for mutations after 32 transfers in their specified heat shock regime. Nucleotide changes are numbered from the start of the NCBI Reference Sequence for the S segment of Φ6 Cystovirus (Accession# NC_003714). Specific mutations engineered, evaluated, and presented in this study are marked with a “Y” in the column “Evaluated for thermostability and fitness?” The column "Pilot lineages" indicates that the mutation was also found in a pilot experiment after 32 days of evolution under constant-temperature heat shock. (The exact heat shock temperature used for these pilot lineages is indicated by the number after "T" in the lineage name, while the number following "R" indicates the replicate number assigned to the population. Mutations found in multiple populations, e.g. populations 1, 2, and 3, are denoted with dashes, e.g., R1-3.)

**S4 Table. Primers used for reverse engineering of** Φ**6 mutants.** The engineered mutation is indicated with upper case in the nucleotide sequence. Where not otherwise stated in Reverse engineering (Methods), primers were prepared according to instructions in the QuikChange II mutagenesis kit. Primers for mutant V109I in P8 were given a standard desalting and used in a T4 ligation reaction; melting temperatures for these primers were calculated using the OligoAnalyzer (Integrated DNA Technologies, http://www.idtdna.com/calc/analyzer) www.idtdna.com/calc/analyzer).

**S1 Fig. Competitive fitness of the ancestral genotype initialized at different frequencies.** Competitive fitness was calculated using Equation 2. The mean competitive fitness is indicated with a dashed line. The competitive fitness values of the ancestral genotype are not significantly correlated with its initial frequency (Pearson’s correlation test, ρ = -0.32, p = 0.30).

**S2 Fig. Changes in percent survival of Sudden populations with and without thermostabilizing mutations**. At each transfer, the survival of each population (Observed percent survival) was compared to the percent survival of the ancestor (Expected percent survival) at 50°C. Lineages are distinguished according to whether at least one mutation evaluated to be thermostabilizing was present in the endpoint population. (Note that, although the entire lineage has been colored for the purpose of visualization, the exact point in time at which the thermostabilizing mutation arose was not evaluated in this study.)

**S1 Text. Effects of additional mutations in the engineered viral mutants on thermostability and relative competitive fitness.**

**S2 Text. Historical sequencing of population Gradual 1.**

